# Altered Fecal Microbiota and Urine Metabolome as Signatures of Soman Poisoning

**DOI:** 10.1101/312660

**Authors:** Aarti Gautam, Derese Getnet, Raina Kumar, Allison Hoke, Amrita K Cheema, Franco Rossetti, Caroline R Schultz, Rasha Hammamieh, Lucille A Lumley, Marti Jett

## Abstract

The experimental pathophysiology of organophosphorus (OP) chemical exposure has been extensively reported. Here, we describe an altered fecal microbiota and urine metabolome that follows intoxication with soman, a lipophilic G class chemical warfare nerve agent. Non-anaesthetized Sprague-Dawley male rats were subcutaneously administered soman at 0.8 - 1.0 of the median lethal dose (LD_50_) and evaluated for signs of toxicity. Animals were stratified based on seizing activity to evaluate effects of soman exposure on fecal bacterial biota and urine metabolites. Soman exposure reshaped fecal bacterial biota by preferentially expanding *Facklamia, Agrobacterium*, *Bilophila*, *Enterobacter*, and *Morganella* genera of the *Firmicutes* and *Proteobacteria* phyla, some of which are known to hydrolyze OPs. However, analogous changes were not observed in the bacterial biota of the ileum, which remained the same irrespective of dose or seizing status of animals after exposure. Interestingly, when considering just the seizing status of animals, we found that the urine metabolome was markedly altered. Leukotriene C4, kynurenic acid, 5-hydroxyindoleacetic acid, norepinephrine, and aldosterone were excreted at much higher rates at 72 hrs in seizing animals, consistent with early multi-organ involvement during soman poisoning. However, at 75 days post soman exposure, bacterial biota stabilized and no differences were observed. These findings demonstrate the feasibility of using the dysbiosis of fecal bacterial biota in combination with urine metabolome alterations as forensic evidence for OP exposure temporally.

**Importance:** The paucity of assays to determine physiologically relevant OP exposure presents an opportunity to explore the use bacterial sentinels in combination with urine to assess changes in the exposed host. Recent advances in technologies and computational approaches have enabled researches to survey large community level changes of gut bacterial biota and metabolomic changes in various biospecimens. Here, we profile combined changes in bacterial biota and urine metabolome due to chemical warfare OP exposure. The significance of our work is to reveal that monitoring bacterial biota and urine metabolites as surrogates of OP exposure in biospecimens suitable for existing clinical laboratory workflows is plausible without the need for the development of new technology, invasive procedures, or complicated analytical approaches. The larger value of such an approach is that any setting with a moderate clinical chemistry and microbiology capability can determine pre-symptomatic exposure to enhance current triage standards in case of mass exposures, refugee movements, humanitarian missions, and training settings once an algorithm has been validated. In the event of “potential” exposures by time or distance, this assay can be further developed to estimate affected radius or time dimension for health monitoring and treatment interventions.

## Background

Despite the serious health threat posed to communities, organic derivatives of phosphorus containing acids have a wide range of applications in modern society (1-3). OP-containing products are in excessive use world-wide for the control of agricultural or household pests. OP-containing pesticides account for almost 38% of all pesticides used across the globe leading to nearly 3 million poisonings, over 200,000 deaths annually, and the contamination of numerous ecosystems (4). In addition, their application as agents of war and terrorism in the form of nerve agents poses a significant threat to both civilians and the warfighter. Exposure to OP leads to various degrees of neurotoxicity due to cholinergic receptor hyperactivity, mediated primarily by the inhibition of acetylcholinesterase (AChE) (5). The excessive accumulation of acetylcholine from the inhibition of AChE leads to severe physiological complications that may manifest both as muscarinic symptoms (e.g. lacrimation, salivation, diarrhea, miosis, and bradycardia) as well as nicotinic symptoms (e.g. tachycardia, hypertension, convulsions, and paralysis of skeletal and respiratory muscles) and can even lead to death (1-3, 6, 7).

Soman, pinacolyl methylphosphonofluoridate or GD (German agent D), is one of the G class nerve agents (volatile agents associated with inhalation toxicity) that inhibits AChE much more rapidly but less specifically than V class nerve agents (viscous agents associated with transdermal toxicity) (2, 6, 8). Whole body autoradiography studies in mice revealed that intravenously administered tritiated-soman (^3^H-soman) spreads within the entire body in less than 5 min (9). High levels of accumulation were noted in lungs, skin, gall-bladder, intestinal lumen, and urine during the first 24 hrs. ^3^H-pinacolyl methylphosphoric acid (^3^H-PMPA), a hydrolyzed acid metabolite of soman, was found to be concentrated in specific organs such as lungs, heart, and kidneys within minutes of ^3^H-soman administration, which reflected the highly reactive (i.e. rapid aging) nature of soman *in vivo* (10). Significant amounts of soman were also detected in red blood cells, a major esterase depot, when compared to the plasma. In addition, these studies revealed that the common route of excretion for PMPA, a major soman metabolite, was via urine and the intestinal lumen content (9, 11). Interestingly, only trace amounts of ^3^H-soman, ^3^H-PMPA, or ^3^H-methylphosphonic acid (hydrolyzed PMPA) were observed in the central nervous system. Current clinical nerve agent exposure assessments are primarily based on overt physiological reactions such as convulsions, loss of consciousness, and salivation for high dose exposures or pupil constriction and, respiratory distress for low dose exposure (12, 13). Recent studies have also demonstrated the feasibility of identifying OP hydrolysis products in hair and nail clippings to verify nerve agent exposure after 30 days (13, 14). Hence, monitoring or verifying sucpected and asymptomatic exposure using minimally invasive and rapid molecular methods will be an ideal approach. Thus, identification of new surrogate biomarkers of toxicity and/or exposure to soman and other OPs is essential both from a clinical and a public health standpoint, especially for triaging population level exposures.

Using an omics approach, we assessed the potential value of changes in fecal biota and urine metabolite as a suitable feature with diagnostic and biosensor utility for OP exposure surveillance and monitoring in a rat model of soman exposure. More importantly, we filled in a knowledge gap of how OP exposure directly or indirectly impacts bacterial communities of the gut and alters the global urine metabolic profile. Applied for more than 20 years in the bioremediation field, specific species from the *Bacteroides* and *Proteobacteria* phylum have been implicated in enhanced biodegradation of OP pesticides. Therefore, exploring the role of the microbiome in a mammalian host’s response to OP is the next logical step (4, 15). Furthermore, recent advances in sequencing technologies have enabled a detailed analysis of structural changes in the gut microbiome revealing the dynamic ecosystem of the bacterial biota and its essential role in health and disease. We also identified urine as a suitable specimen type for investigation in this study design because urine consists of numerous metabolites as outputs from multiple pathways and provides a snapshot of both local and systemic physiological changes (16). With this in mind, we focused our efforts on exploring and describing soman-induced dysbiosis of the gut microbiota and alterations in urine metabolome from a systems level analysis.

## Results

### Clinical manifestation of soman insult

To establish soman-induced toxicity with and without seizure, Sprague-Dawley rats were subcutaneously injected with saline or 0.8 or 1 LD_50_ equivalent of soman. To reduce mortality, the animals given 1 LD_50_ were also administered atropine sulfate and HI-6 one minute after exposure. Rats that developed seizures at 1 LD_50_ were additionally given diazepam to control seizing. Approximately 42% of the animals exposed to soman (0.8 or 1.0 LD_50_) experienced seizure irrespective of dose or the medical treatment regimen administered. As expected, control rats did not experience seizure from administration of the vehicle. Seizing animals experienced a notable weight loss and displayed increased activity in the days immediately after soman poisoning (Fig. 1a and Fig. S1a). Body temperature was not altered between seizing and non-seizing groups (Fig. S1b). The Racine scale score, a quantitative assessment of seizure-related activities such as degrees of tremors, convulsions, and seizures, was significantly higher in seizing subjects as compared to non-seizing subjects, as expected (Fig. 1b)(17-19). Based on EEG activity, body weight, and Racine score, we broadly categorized our analysis groups into a non-seizing group (no seizure, n= 13) or a seizing group (exposure seizure or sustained seizure, n=10). We also further subdivided cohorts based on dose because of seizure differences (Fig. 1c and 1d). Fecal matter, urine, and tissues were harvested from animals to examine the gut microbiota and urine metabolic changes due to the soman insult.

**Figure 1.**
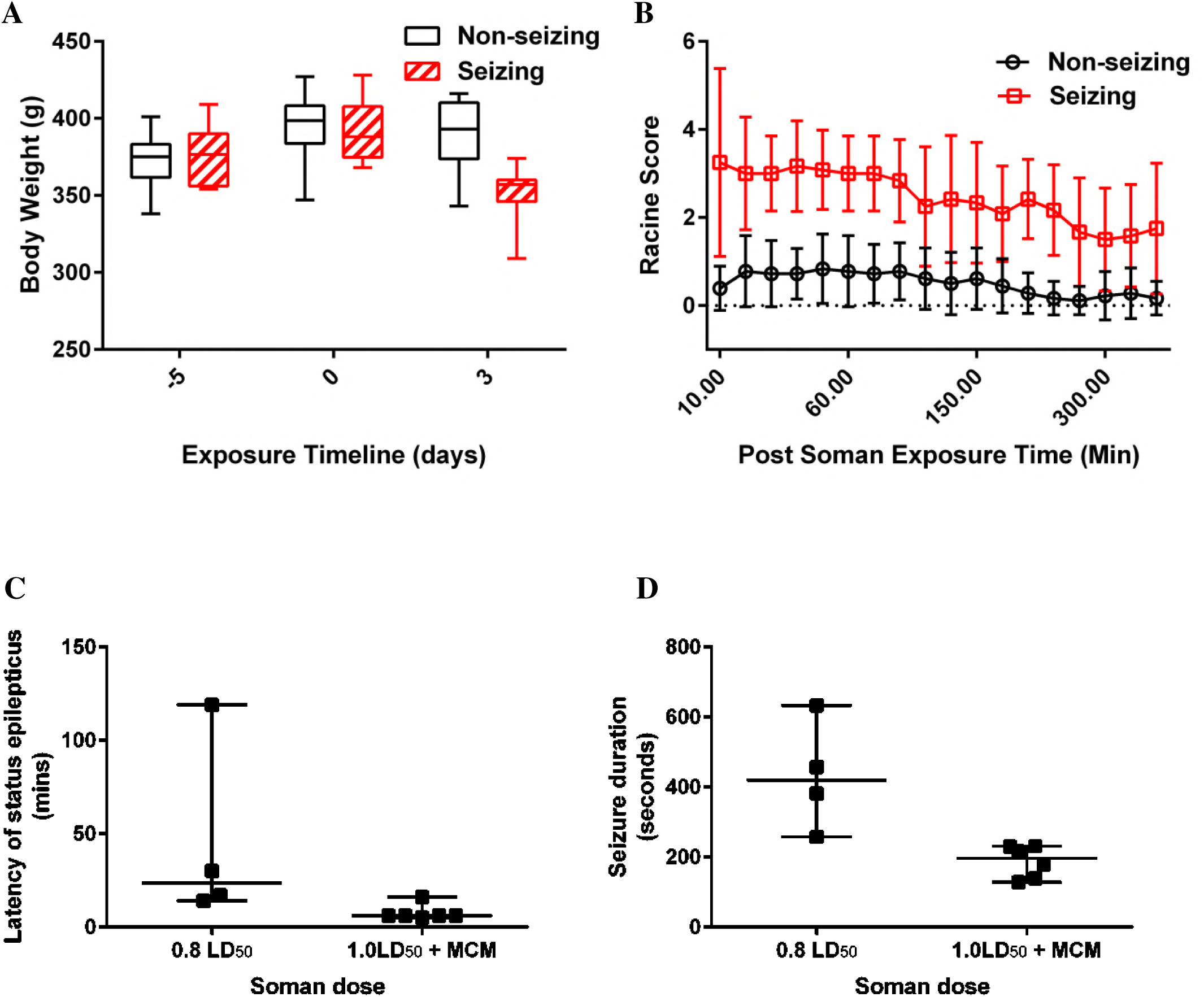
Clinical manifestation of soman exposure. Panel a represents trends in body weight changes between non-seizing and seizing rats. Panel b represents Racine Score differences between non-seizing and seizing rats. Panel c and d represents seizure activities (latency to seize or seizure duration during 72hr monitoring) of rats exposed to 0.8 LD_50_ of soman without treatment or 1.0 LD_50_ of soman and treated with medical counter measure (MCM) a minute after exposure.

### Taxonomic changes to soman exposure

To assess the effect of soman exposure on the bacterial biota, we sequenced the hypervariable regions of the 16S ribosomal RNA from the feces and ileum of all animals at 72 hrs. Collecting and sequencing specimens from individual animals enabled us to assess the effect of dose- or seizing status- driven changes in individual biota. By measuring multiple alpha diversity estimates between dose (0.8 LD_50_ vs 1.0 LD_50_) and seizing status (non-seizing vs seizing), we found statistically significant (*p<*0.05) increased distances indicating altered diversity in the fecal bacterial biota of seizing animals at both 0.8 LD_50_ and 1.0 LD_50_ (with a pronounced effect in seizing animals of the 0.8 LD_50_ exposure) that was markedly absent in control or non-seizing groups (Fig. S2a). When comparing alpha diversity distance between the ileum specimens of control and soman exposed animals, however, the distances were much smaller indicating higher degrees of similarity among taxa within the various groups, irrespective of dose or seizing status (Figure S2b). The bacterial biota of the ileum was not altered by exposure to soman while the fecal bacterial biota was substantially altered by all measures of diversity.

In order to investigate if organism abundance accounted for the alpha diversity differences observed between the fecal and ileum bacterial biota’s response to soman insult, we rank ordered all phyla identified in the study based on relative abundance (data not shown). To our surprise, no differences were observed between the fecal and ileum bacterial phyla abundance distribution. In both fecal and ileum specimens, *Firmicutes* and *Bacteroidetes* accounted for the most abundant phyla as expected followed by *Verrucomicrobia*, *Tenericutes*, and *Actinobacteria*. However, within the ileum the relative abundances of all the phyla were closely distributed around 10% while the phyla within the fecal specimens displayed a wide range of distribution from 6% to 18% (data not shown). To further understand if soman exposure-driven bacterial biota differences were attributable to dose or seizure individually, we measured beta diversity across different specimens (fecal or ileum) and visualized the output using principal coordinate analysis (PCoA) (Fig. S3). This PCoA, however, was unable to distinguish any of the groups into significant clusters based on dose or seizing status in either the ileum or fecal specimens.

### Effect of soman exposure on microbiota compositions

We next examined the microbial communities of the feces and ileum of each subject and enumerated each phylum to decipher where microbial diversity was substantially altered by exposure to soman, based on dose or seizure status. Several structural changes were observed in the fecal microbiota compositions correlating to dose or seizure status of animals (Fig. 2). We found that the fecal *TM7* phylum relative abundance was substantially reduced in response to increasing dose of soman and seizure status of animals, concurrently. Conversely, fecal *Proteobacteria* and *Cyanobacteria* expanded in response to increased soman dose insult and seizing of animals(Fig S4). The relative abundance of *Actinobacteria*, *Firmicutes*, *Tenericutes*, and *Verrucomicrobia* phyla, however, was unchanged in response to soman dose or seizing status of animals. The *Bacteroidetes* phyla showed a positive trend of expansion, although not statistically significant. Similar microbial community composition analyses were also completed for ileum specimen collected from each individual subject to identify if anything was masked during the initial analyses (Fig. S5). Consistent with previous findings in the ileum, no significant structural bacterial biota changes were observed in any subjects in response to dose or seizing status.

**Figure 2.**
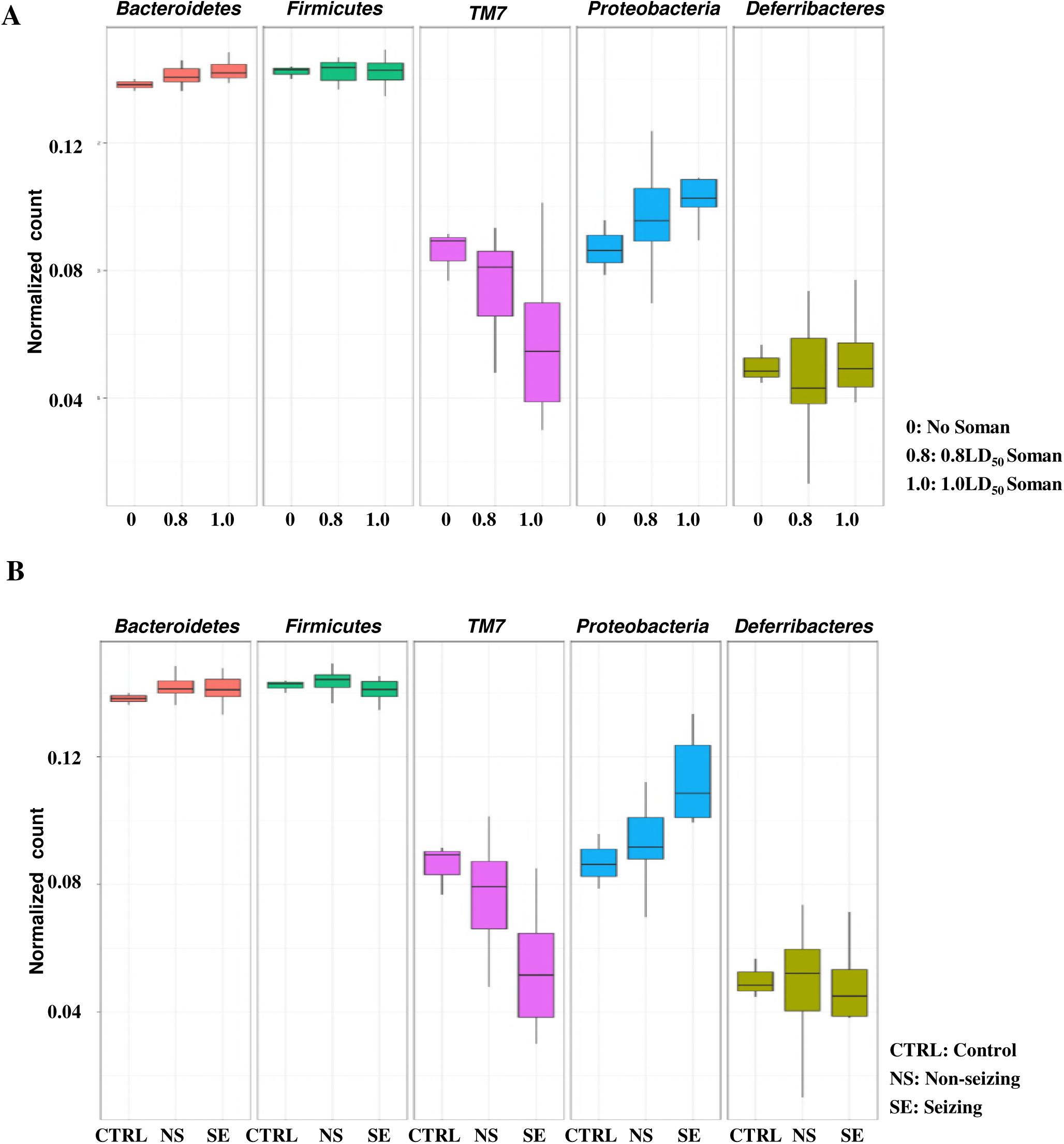
Altered fecal microbiota composition post soman exposure. Trends in relative abundance of fecal bacterial biota based on dose (Panel **a)** and seizing status (Panel **b**) of animals

To measure the bacterial biota diversity and relative abundance (with respect to quantitative and qualitative analysis), we further looked into the taxonomic breakdown of each phylum into cohesive genetic clusters, OTUs (operational taxonomic units), used to construct the phylum level data. Distinct bacterial populations were observed at the genus level of *Firmicutes* and *Proteobacteria* in the fecal specimens of soman exposed animals (Fig. 3). The *Facklamia* genus of the *Aerococcaceae* family was only observed in soman exposed rats. This *Firmicutes* genus was also detected only in the fecal specimens of soman-exposed rats and not in the ileum tissues of these cohorts. *Agrobacterium*, *Bilophila*, *Enterobacter*, and *Morganella* genera of the *Proteobacteria* phylum exhibited expansion in the soman exposed rats similar to that observed for the *Facklamia* genus. Interestingly, the *Agrobacterium* genus was primarily identified in the lower dose (0.8 LD_50_) soman exposure group while *Bilophila*, *Enterobacter*, and *Morganella* genera were detected in all soman doses irrespective of seizing status of animals (Fig S6). Furthermore, several unclassified genus clusters were observed for *Alcaligenaceae*, *Comamonadaceae* and *Enterobacteriaceae* families of the *Burkholderiales* order, primarily in the seizing high soman dose exposure animals, and these genus clusters were not observed in the control group or the low soman dose exposure group. All difference in bacterial biota composition here were only observed at the 72 hr time point and neglible at 75 days post exposure (data not shown).

**Figure 3.**
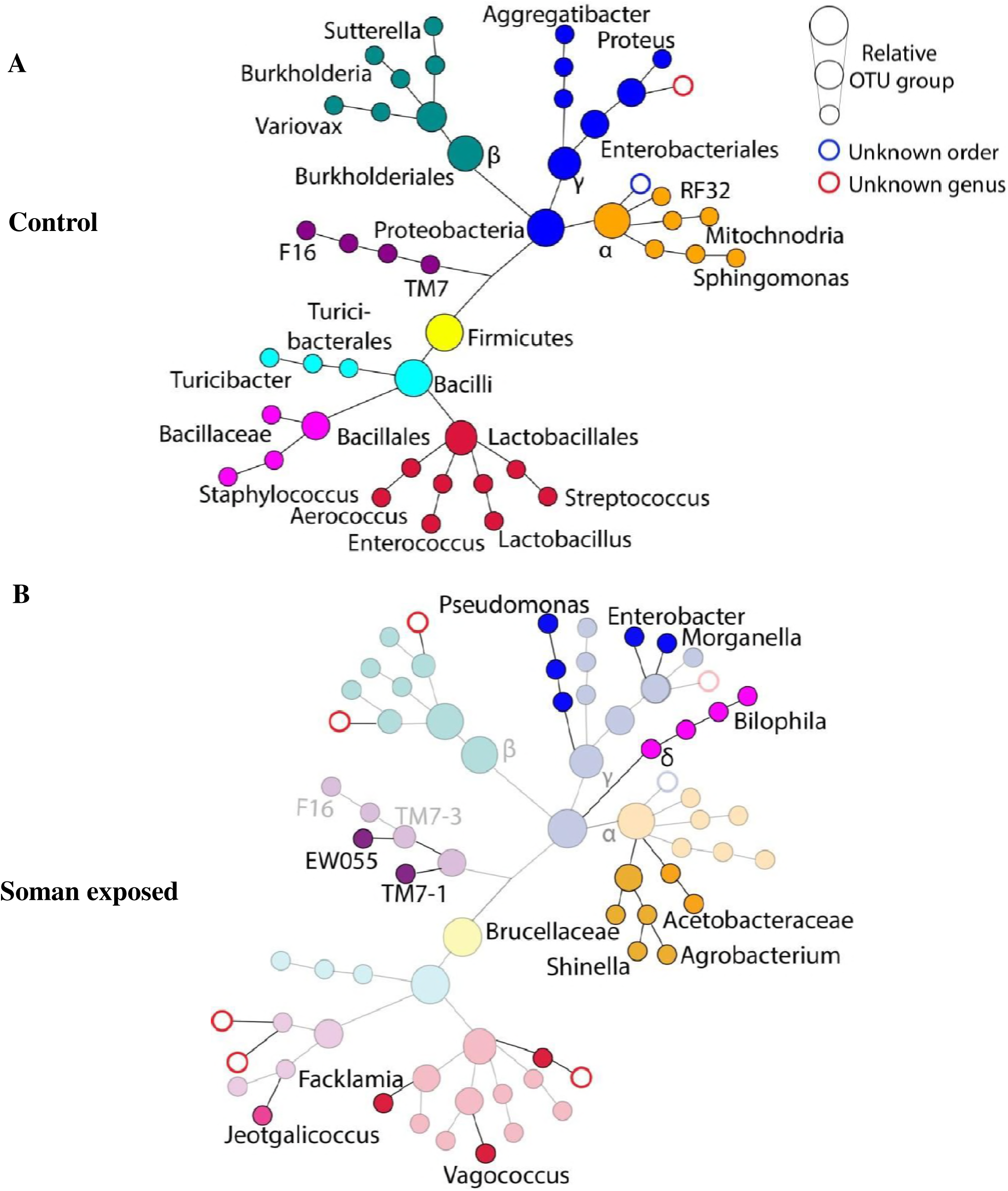
SPADE-like analysis showing taxonomic changes in *Proteobacteria* and *Firmicutes* (*Bacilli* only) phyla. Panel **a** represents average fecal bacterial biota detected for the two phyla in soman unexposed control animals. Panel **b** shows representative average of expanded bacterial biota detected after exposure (solid circles) compared to bacterial biota of unexposed control (faded circles). Undetected bacterial biotas are represented by missing circles. Distance between circles is assigned arbitrarily.

### Soman exposure alters the urine metabolome

The untargeted metabolomic profiling of biological substrates provides direct and simultaneous measurements of biochemical outputs that make up a given phenotype (20). Comparative metabolomics profiling of urine specimens was performed to assess soman exposure associated metabolic alterations at 72 hrs post exposure, for seizing and non-seizing animals. XCMS was used to pre-process metabolomics data to generate an analysis matrix of mass-over-charge, retention time, fold change, and *p* value for over 500 unique metabolite peaks identified from both positive and negative mode analyses. Volcano plots from Metaboanalyst v3.0. were used to visualize and enrich for metabolites with ≥ 2 fold change at *p*<0.05 (Fig. S7). Furthermore, PCA was used to visualize if seizing and non-seizing groups clustered separately (Fig S8). At the 72 hr time point, we found >200 metabolite peaks with >2.0 fold change, of which 9 analytes had >20 fold change in seizing rats rather than non-seizing rats. Many of these metabolites with >20- fold change appear to be either food-drived metabolites such as methylmaysin and acetylpicropolin, or cresol derivatives. We also found a large fraction of modified amino acids (such as acetylated, methylated, or formylated) common urine solutes along with many phenyl and indole compounds. We used an in-house algorithm to identify bacterial metabolites that are known to be detected in urine. Hence, we identified routine urine metabolites associated with microbial origin such as p-cresol, D-alanine, and phenylacetic acid. Interestingly, these metabolites were detected at a significantly higher rate (>5 fold change) in the urine of seizing rats than non-seizing rats. The tryptophan catabolism by product kynurenic acid, the inflammatory lipid leukotriene C4, and the neurotransmitter norepinephrine were also secreted in the urine at a significantly higher rate in seizing animals (Table 1)(21). However, urinary citric acids leveles remained unaltered between seizing and non-seizing animals, suggesting that the presence of physiological indicators of physiological dysregulation (acidemia and inflammation) in the urine of soman-exposed animals was not primarily the result of kidney injury.

**Table 1.**
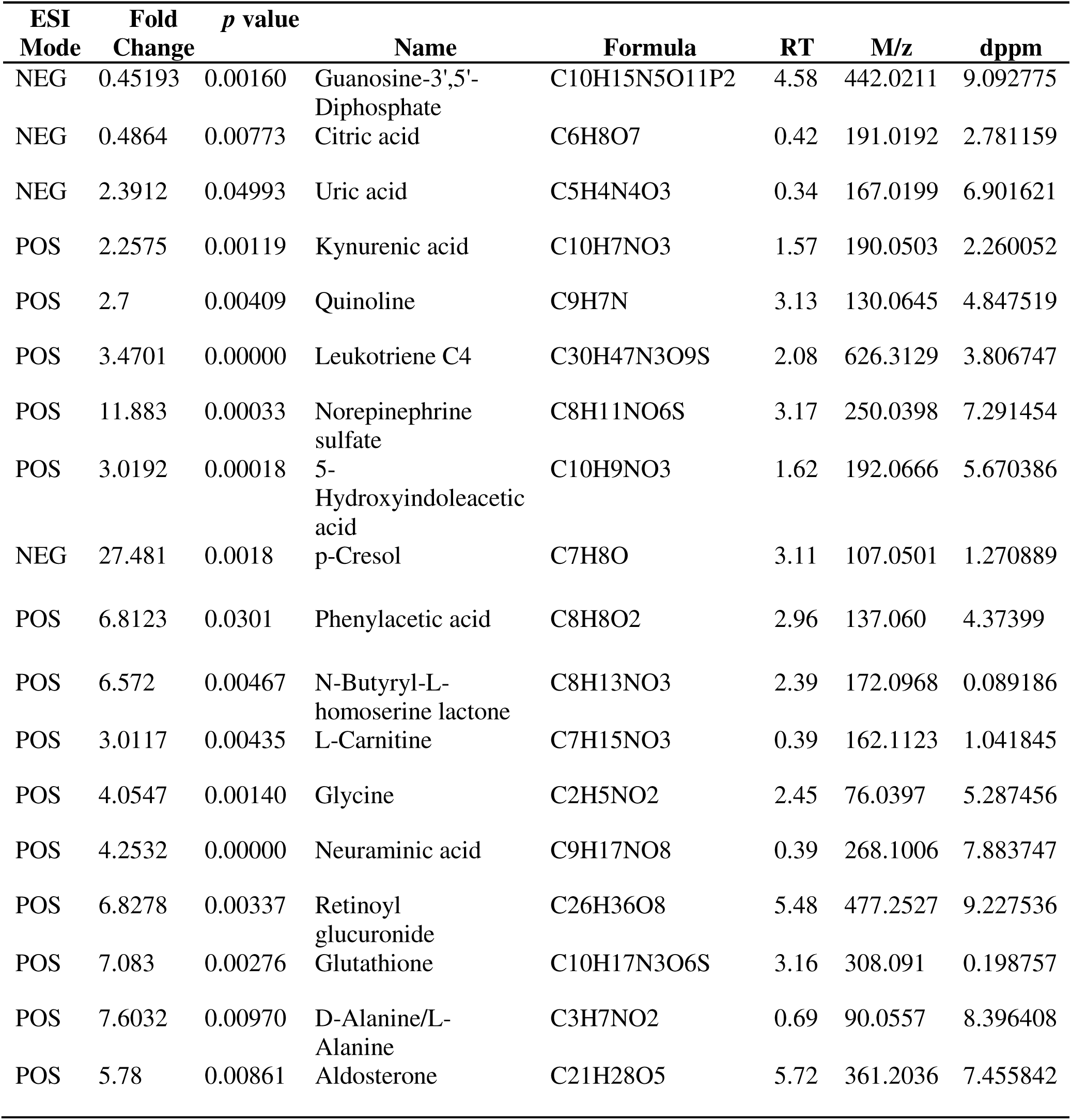
Summary of soman-exposure altered urine solute

To gain insights into biological networks perturbed by soman in seizing rats, we annotated all molecular networks associated with candidate metabolites identified at the 72 hrs time point in an unbiased manner using publically available databases. Consistent with the neurotoxicity pathology of soman assault, large proportions of the metabolites in seizing animals’ urine were primarily related to nervous system signaling activity representing norepinephrine and serotonergic pathway products (data not shown). We next narrowed down our pathway annotation by only considering best peak-matched metabolites (such as highest mass accuracy (lowest dppm) and frequent database hits.) and manually removing unlikely identifications, such as drug action pathways, to restrict off-target hits in our annotation algorithm. This careful focus of the data input revealed key molecular networks (catabolic processes) reflective of products associated with kidney, canonical central nervous system (CNS) inflammation, amino acid and lipid metabolism, and vitamin absorption (Fig. 4). To increase the confidence in this analysis, a select set of metabolites were validated in pooled specimen from seizing and non-seizing subjects.

**Figure 4.**
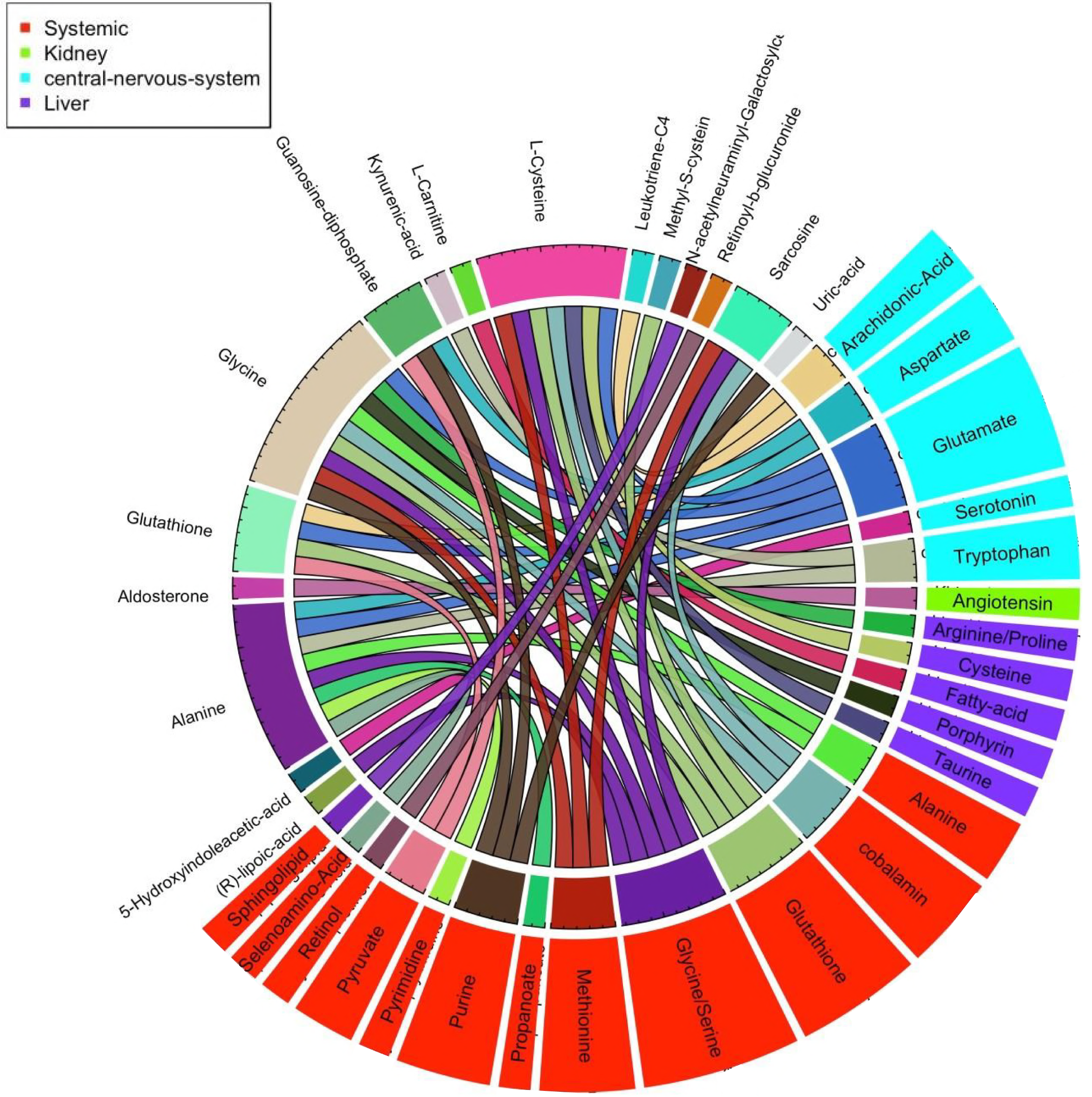
Snapshot of altered urine metabolites, affected candidate site, and pathways in non-seizing and seizing rats. Circos image represents urine solutes significantly altered (*p<0.05*) in seizing animals (outer ring labels on white background) and associated metabolic pathways and sites (outer ring colored boxes).

## Discussion

Here, we employed a high throughput systems approach to survey changes in the gut bacterial biota and urine metabolites and to explore new areas of potential diagnostic markers for OP exposure and toxicity in noninvasively collected specimens. We initially noted that seizing subjects were significantly different than controls and non-seizing subjects based on multiple parameters, including alpha diversity measures. However, composition analysis indicated that the *TM7* and *Cyanobacteria* phyla were probably the main drivers of the large observed alpha diversity differences while the *Proteobacteria* phylum was a modest contributor of these observed differences. Unfortunately, *Cyanobacteria* represent organisms such as chloroplasts and the *TM7* phylum is not well understood. Due to limited knowledge, we concluded that these two phyla are insufficient for drawing meaningful conclusions for the observed biological differences at this moment. Thus, we focused our efforts to closely examine the genera compositions that make up the various phyla identified in our study especially *Proteobacteria*. To this end, we noted the presence of *Facklamia* genus, a *Firmicutes*, only in soman exposure feces and not in the ileum while the *Agrobacterium*, *Bilophila*, *Enterobacter*, and *Morganella* communities of *Proteobacteria* were temporally altered by the soman insult independent of seizing status or dose.

In our system, the dysbiosis of these microbial populations represents the interplay between resistance and resilience to soman-induced physiological and ecological stress temporaly. Hence, the selective expansion or contraction of a given genus is either to fill in the space created by the decrease in the soman-stress-susceptible communities, change in fermentative environment of the gut, or a result of exploiting metabolic advantages of soman hydrolysis, which is documented for some bacteria in the environment. It is important to point out that many of the organophosphate-degrading genes (*opd*) implicated in the enhanced degradation of OPs are located on mobile elements known to be transferred between organisms (22). In addition, a majority of these genes exhibit a broad specificity and activity against OPs by either directly hydrolyzing the phosphoester bonds in organophosphorus compounds or by further degradation of methylphosphonate-esters to reduce toxicity or reactivation of OP (4, 22). Such genes have been isolated from select species of *Flavobacterium, Pseudomonas, Alteromonas Burkholderia, Bacillus, Alcaligense, Enterobacter, and Agrobacterium* genera to name a few. Thus far, these genes and organisms have not been implicated in *in vivo* bioremediation in mammalian hosts to date. In this study, however, some organisms primarily within *Proteobacteria* phylum such as *Agrobacterium*, *Enterobacter*, and *Morganella* genera, known to be involved in the enhanced degradation of OPs in the environment, were also detected here as having expanded temporally in response to the soman insult *in vivo*.

The *Facklamia* genus was detected in the feces but not in the ileum tissue of rats exposed to soman. However, this expansion was masked by the dominant nature of the *Firmicutes* phylum whose overall relative abundance did not change in this exposure study. *Facklamia* are facultative anaerobic alpha-hemolytic, Gram-positive cocci that are part of the normal fecal flora often mistaken as viridans streptococci (23-25). This genus is hardly associated with invasive disease, but it is known to be occasionally isolated from specimens of urinary tract infections and chorioamnionitis infection. To date, there are six species of *Facklamia* of which the first four have been isolated from humans: *F. hominis, F. ignava, F. sourekii, F. languid, F. tabacinasalis sp nov,* and *F. miroungae sp nov.* The pathogenic potential of these species is unclear. Similarly, limited data is available regarding their metabolic functions specifically hydrolysis, reduction, or esterification of xenobiotics within the gut, although such functions have been described for other *Bacilli* spp from the *Firmicutes* phylum.

*Agrobacterium*, an *Alphaproteobacteria*, currently renamed as *Rhizobium*, is an aerobic, oxidase-positive, Gram-negative bacillus isolated from the soil environment and known to cause plant tumors (26, 27). They are opportunistic human pathogens primarily associated with outbreaks of sterile-site and catheter-associated infections. Interestingly, *Rhizobium radiobacter* is known to directly hydrolyze a wide range of organophosphate insecticides through OPDA, a unique OP-degrading *Agrobacterium* (*opdA)* chromosomal gene (4, 15). In our study, the *Rhizobium* genus appears to have selectively expanded primarily in the low dose soman insult group (0.8LD_50_) but is absent in animals at the higher dose. This observation, however, is confounded by treatment regimens provided to animals at 1.0 LD_50,_ for which the relevant controls were not within the scope of this study design and must be investigated in a future study. Currently, there is limited information about which species of *Rhizobium* resides in the gut flora of mammalians as a routine commensal, which will be possible to identify with whole genome sequencing approach. Its presence here suggests that the introduction of *Rhizobium* to the rat colony may have occurred through food intake considering its association with plants and plant roots (15, 26). Furthermore, the functional contributions of *Rhizobium* to the gut need to be closely examined to establish any functional role within the gut microbiota.

The *Bilophila* genus is the only known *Deltaproteobacteria*. This genus is an obligate anaerobic Gram-negative bacillus comprised of only a single species, *Bilophila wadsworthia*, which is associated with normal human fecal and vaginal flora (28). This bacterium is an opportunistic pathogen commonly isolated from anaerobic infections of the abdomen, including appendicitis. *B. wadsworthia* is known to thrive on taurine and H_2_ production around highly fermentative sites. Unlike the *Rhizobium* genus, *B. wadsworthia* was highly enriched and detected in all soman exposure cohorts, indicating the creation of a highly anaerobic environment in the gut, altering the fermentative condition that promoted the expansion of this organism.

*Morganella*, an *Enterobacteriaceae* and a *Gammaproteobacteria*, is an aerobic Gram-negative bacillus principally isolated as normal fecal flora (29). This genus contains two species: *Morganella morganii* and *Morganella sibonii*. Unlike *Bilophila wadsworthia*, *Morganella* was primarily associated with seizing subjects of both high and low soman exposure, as opposed to the non-seizing subjects exposed to soman or the control animals. Unfortunately, very little is known about the biochemical conversion roles of the *Morganella* genus in general and its role in xenobiotics conversion. Another *Enterobacteriaceae* tribe that expanded in response to soman exposure is *Enterobacter*, a Gram-negative bacillus that is a catalase-positive facultative anaerobe. Many of the 14 *Enterobacter* species are found in the intestinal track and skin of humans and animals as well as in the environment. This genus is primarily known for being an opportunistic pathogen in humans. Several environmental *Enterobacters,* such as *E*. *asburiae*, have been implicated in the biodegradation of OPs through a known plasmid-carried *opd* gene (4, 30). Hence, mobile elements within this genus may provide the metabolic advantage that contributes to expansion of *Enterobacter spp* in this setting.

Seizures that accompany soman intoxication lead to profound brain damage through excitotoxic cell death of neurons, accompanied by neuroinflammation (31). Since we observed significant correlation between the various physiological parameters (body weight, activity, and Racine score) with seizing or non-seizing status, we broadly grouped exposed animals accordingly to identify relevant urinary solutes that can serve as phenotypic markers of consequential soman exposure. Therefore, we were able to increase our signal-to-noise ratio and enrich multiple microbial and mammalian co-metabolites and their respective pathways that correlated with symptomatic soman exposure. We detected elevated levels (>2-fold change) of tryptophan catabolism, namely kynurenic acid and quinoline, in the urine of seizing animals. Kynurenic acid is a putative neuroprotective metabolite as an antagonist of N-methyl-D-aspartate (NMDA) receptors (21, 32, 33). Interestingly, we also detected equivalent, elevated levels of quinoline, a reduced version of quinolinic acid which is an agonist of NMDA receptors and a neurotoxic metabolite of tryptophan. These findings, however, clearly indicate an overall increased tryptophan catabolism in the system during soman intoxication in seizing animals, presumably driven by inflammation in the brain, which is a known inducer of the tryptophan degrading indoleamine 2,3 dioxygenase (IDO) (32, 33). This notion of the inflammatory neuropathology of soman exposure was further supported by the detection of increased systemic levels of the eicosanoid inflammatory mediator leukotriene C_4_ (LTC_4_) in seizing animals’ urine. LTC_4_ is one of the cysteinyl leukotrienes (cys-LTs) generated from the enzymatic oxidation of arachidonic acid, a fatty acid released from neuronal membrane glycerophospholipids during a secondary phase of brain injury (via a cascade of physiological reactions to primary injury) (34, 35). Cys-LTs including LTC_4_ have many biological activities, including superficially increasing plasma extravasation and formation of inflammatory edema in tissues. In an experimental stroke brain injury model, cys-LTs induced blood brain barrier disruption and brain edema to exacerbate CNS injury (34). Urinary excretion of systemically elevated cys-LTs is a feature of numerous heterogeneous disorders (35). In our experimental system, the primary stimulant soman is known to cause severe neuropathology specifically inflammation of the CNS, and altered immune cell function in seizing animals (31). In this context, the increased levels of the cys-LT LTC_4_ can be attributed to and an indication of an inflammatory CNS neuropathology that was specifically enhanced in seizing animals as compared to non-seizing animals.

Numerous chemical changes have been observed in different parts of the brain during soman-induced seizures (36). These changes appear to occur in several waves. For instance, immediately after acute soman intoxication and seizure onset, high levels of acetylcholine accumulation in the CNS are followed by a decreased level of norepinephrine (NE) in the brain along with aspartate and glutamate (GLU) (i.e. excitatory amino acid (EAA) neurotransmitters) while dopamine and its metabolites are significantly increased, along with gamma-aminobutyric acid (GABA). After neuronal damage, however, EAAs like GLU increase in concentration and, in addition, there is an increase in the release of NE (36, 37). Furthermore, elevated levels of serotonin (5-HT) and its’ metabolite 5-hydroxyindolacetic acid (5-HIAA) accompany the onset and sustainment of seizures (31). Overall, these studies highlight the increased turnover of multiple neurotransmitters and their metabolites, leading to and involved in the cascade of long-term seizure pathology. Consistent with this notion, we observed increased levels of NE (> 11- fold change) and the serotonin metabolite 5-HIAA in the urine metabolite profile of seizing animals.

Aldosterone is an essential mineralocorticoid hormone directly involved in the regulation of sodium absorption and potassium excretion in the kidney, salivary glands, sweat glands, and colon (38). Elevated levels of aldosterone are known to alter glomerular structure and function via pro-oxidative and pro-fibrotic changes (39). Hence, aldosterone increases glomerular permeability to albumin, leading to increased protein urinary excretion. We found that urinary excretion of aldosterone is more than 5 times higher in seizing rats than non-seizing rats, indicating that symptomatic soman exposure causes proteinuria and increased electrolyte excretion. Furthermore, in this context, aldosterone driven nephropathy and renal damage of modest effect are possible secondary features of soman insult. However, the experimental design here did not include analysis of renal function or urine composition, specifically protein and electrolyte excretions.

Xenobiotic metabolites of microbial origin have been previously detected in urine (40, 41). In our study, we identified a few of the most common microbial urinary metabolites, p-cresol and phenylacetic acid (42). Both of these metabolites are a result of tyrosine catabolism by colon microbes and sulfated by the colonic epithelium, resulting in reabsorption back into the host. High levels of p-cresol sulfate are correlated with cardiovascular diseases as well as the mortality of chronic kidney disease patients (40). Furthermore, the phenolic features of such compounds are speculated to mimic neurotransmitters and interfere with the blood-brain barrier(40). Consistent with our finding of dysbiosis of the gut bacterial biota, the soman exposed animals, in our study had a significant increase in p-cresol (27x) and phenylacetic acid (7x) in their urine as compared to non-seizing animals. However, we did not obtain enough 16S sequencing resolution for species-level information in our efforts to identify organisms, especially those associated with tyrosine catabolism such as *Clostridium difficile* (40).

In experimental models of soman exposure, some subjects may develop increasing seizing activity that progresses to *status epilepticus* (*SE*) with profound brain pathology if seizures persist for more than 30 minutes, while other subjects remain free of seizure activity and do not develop brain pathology (31, 36). Seizures, in this context, are classified into 3 phases: cholinergic phase - early seizure due to acute toxic event (<5 min post-exposure), transitional phase - mixed cholinergic/noncholinergic seizure (5–40 min post-exposure), and non-cholinergic phase neuronal damage-mediated seizure (>40 min post-exposure). In this light, our experimental system has some limitations in which administration of 0.8 LD_50_ or 1.0 LD_50_ with medical treatment caused *SE* unevenly limiting our analysis to broad categories. In addition, limited time points and dose schedules did not exclude the possibility of catching early trends in our current observations, which we plan to address in the next round of experiments. However, this has enabled us to perform blinded analysis on all possible combinations and identify significant differences computationally to correlate findings to physiological data available on each cohort.

Although this was a feasibility study that applied untargeted technologies, we were able to enrich for candidate diagnostic markers both from the bacterial biota and urine solutes studies and also identify organisms with potential role as forensic signatures of exposure. Since the gut conditions that favor *B. wadsworthia* are known and it is the only species in its class, we decided to validate if the relative OTU using a PCR assay targeting the 16S on pooled fecal DNA extracts (Fig. S9). We observed that *B. wadsworthia* levels were altered especially in seizing animals with high dose exposure, but were faintly expressed in controls. As an indirect signature of OP exposure resulting from the general alteration of the fermentative conditions of the gut, *B. wadsworthia* represents one of the many candidates that need to be further investigated in humans. In our context, the detection of expanded *B. wadsworthia* combined with urine metabolite in a quantitative and qualitative matrix could enable a contextual surrogate marker for OP exposure driven pathology. Specifically, urine metabolites such as aldosterone, leukotriene C_4,_ norepinephrine, and 5-hydroxyindolacetic acid, which could be assayed in clinical chemistry workflow, represent a viable set of markers to be closely studied in non-seizing soman exposed subjects in temporally collected specimens. Unfortunately, our current experimental design was not able to accommodate this line of inquiry with respect to early specimen collection for examinable trends. However, we gained insight into a potential forensic signature role of some of the alimentary canal organisms including previously described classes of organisms with bioremediation function. We had identified multiple genera known in to be involved in enhanced degradation to include *Enterobacter* and *Rhizobium*, known carriers of *opd* genes. Here, we were able to amplify PCR products from pooled fecal DNA extracts indicating the expected gene templates were present in the bacterial biota (Fig. S9). However, we have not had any evidence for either a specific species carrying the gene templates of *opd* genes or data supporting if the genes are expressed (no RNA or protein). With additional resources and better targeted technology, we plan to pursue a more detailed analysis on a large cohort of animals.

Current biomarker efforts with respect to nerve agent/OP exposure fall into two major categories: free OP metabolites or protein adducts in biofluids. Many of these efforts require specialized instrumentation, highly skilled technicians, and complex extraction methods of biofluids. Our unbiased systems level approach to analyze a snapshot of urinary solutes and bacterial biota of the gut in soman- exposed animals opens a new avenue for functional markers of soman, other nerve agents, or OP exposure that could be easily adopted to current clinical pathology workflows. The identification of the expansion of OP/derivative catabolizing microbial communities in the gut of rats, such as *Agrobacterium*, *Enterobacter spp, morganella* as well as significantly elevated uremic metabolites, such as aldosterone, leukotriene C_4,_ norepinephrine, and 5-hydroxyindolacetic acid, fill in knowledge gaps of secondary physiological effects in OP exposure pathology. *B. wadsworthia* represents one contextual indirect biomarker that needs further investigation in combination with assessment of the fermentative activity in the gut. Bacterial biota difference were also only relevant at the 72 hrs post exposure and not 75 days after exposure indicating the importance of timing for sampling. The alterations of the urine metabolites observed also provided a list of candidate solutes that require a targeted metabolomics study in the future. Based on these findings, one can envision simpler practical technologies, a coorelation matrix, and decision algorithm to enahnce current clinical microbiology and urine chemistry workflows for diagnostics purposes especially pre-symptomatic phase, determining the radius of impact during mass exposure, screen training sites, and movements of refugees across time dimension and in non-invasive manner by simply leveraging readily available resources in clinical pathology.

## Methods and Meterials

### Animals

Male Sprague-Dawley rats (250-300 g; Charles River Laboratories (Kingston, NY)) were individually housed on a 12:12 hrs light cycle with ad libitum access to food and water. Rats were weighed daily. The experimental protocol was approved by the Institutional Animal Care and Use Committee at the United States Army Medical Research Institute of Chemical Defense(IACUC U-908), and all procedures were conducted in accordance with the principles stated in the Guide for the Care and Use of Laboratory Animals and the Animal Welfare Act of 1966 (P.L. 89–544), as amended.

### Surgery and EEG recording

Rats were surgically implanted with a subcutaneous transmitter (F-40EET; Data Sciences International, Inc. (DSI; St. Paul, MN)) as described in Schultz et al. (2014), to record bi-hemispheric cortical EEG waveform activity as well as body temperature and activity throughout the duration of the experiment. Surgery was conducted under isoflurane (3 - 4% induction; 1.5 - 3% maintenance) and rats received buprenorphine (0.03 mg/kg, sc; Reckitt Benckiser Pharmaceuticals, Inc, Richmond, VA) immediately after full recovery from anesthesia. Rats recovered from surgery for 7-14 days prior to soman -exposure. RPC-1 physiotel receivers (DSI) were placed under the rats’ home cage for EEG acquisition (24 hrs/day) with baseline recordings made at least 24 hrs prior to exposure. Data were digitized at 250 Hz and recorded using Dataquest ART 4.1 (Acquisition software; DSI).

### Soman Exposures

Rats were exposed 0.8 or 1.0 LD_50_ soman (0.5 mL/kg, using 157.4 and 196.8 μg/ml respectively) or saline (control) and evaluated for seizure activity. Soman LD_50_ is 98.4 μg/ml (43). Soman was obtained from the Edgewood Chemical Biological Center (Aberdeen Proving Ground, MD). To promote survival, rats that received 1.0 mg/kg soman were treated with medical countermeasures (MCM; an admix of 2 mg/kg atropine sulfate (ATS, Sigma-Aldrich Chemical Company, St. Louis, MO, USA) and 93.6 mg/kg HI-6 (0.5 ml/kg, im, Starkes Associates, Buffalo, NY, USA)) at 1 min after exposure and rats that developed seizures were treated with 10 mg/kg diazepam (DZP; 2 ml/kg, sc, Hospira Inc., Lake Forest, Illinois, USA) at 30 min after seizure onset (with average seizure onset of 8 min).

### Behavioral seizure

Behavioral seizures were scored using a modified Racine scale (17-19): stage 1: mastication, tongue fasciculation, oral tonus; stage 2, head tremors, head bobs; stage 3, limb clonus or tonus, body tremor; stage 4, rearing with forelimb clonus; and stage 5, rearing and falling with generalized convulsions. For analysis, rats received a score corresponding to the maximum stage reached per time interval. Observations were made continuously for up to 5 hrs after exposure to soman.

### EEG analysis

Full-power spectral analysis of EEG, identification of epileptiform activity, and other EEG anomalies were analyzed according to the methods described previously (44). EEG recorded seizures were confirmed through visual screening and characterized by sustained frequencies and the value of the most prominent frequencies in Hz (e.g.: the highest power calculated by MatLab program (www.mathworks.com) in μV2/Hz). Some soman -exposed rats developed seizure activity (SE) with seizure onset at ~ 8 min for those exposed to 1.0 LD_50_, while others did not develop seizures.

### Sample collection

All surfaces and tools, including the guillotine, collection foils, sample tubes, saline bottle, and syringes, were sprayed with RNase AWAY^®^. Animals were administered Fatal-Plus^®^ (sodium pentobarbital) and once fully anesthetized the rats were euthanized using a guillotine. Urine was collected directly from the bladder using a syringe with 21 gauge needle, saved in RNase-free microfuge tubes, and flash frozen in liquid nitrogen. The ileum was removed and flash frozen. Digesta was flushed out by rinsing with sterile saline during sample processing. Organs were collected in foils and flash frozen in liquid nitrogen. Fresh fecal pellets were collected during rat handling and stored at −80°C for processing.

### DNA extraction

Samples were kept cold at all time before the extraction. The Ileum tissues were weighed and homogenized in 50 mM Tris-HCl (Lonza, Walkersville, MD, USA) with 2 nM EDTA Solution (Lonza) using the BeadBeater (Bio Spec Products, Inc., Bartlesville, OK, USA), the DNA extraction was carried out using the QIAGEN DNeasy Blood and Tissue Kit (QIAGEN Inc., Germantown, MD, USA), and RNA was extracted using TRIzol Reagent (Invitrogen, Life Technologies, Grand Island, NY, USA) in conjunction with the QIAGEN miRNeasy Mini Kit (QIAGEN). The fecal samples were weighed and RNA was extracted using MoBio PowerSoil Total RNA Isolation Kit (MO BIO Laboratories, Inc, Carlsbad, CA, USA) and DNA was extracted using the MoBio PowerSoil DNA Elution Accessory Kit. The extracted DNA was used for PCR and sequencing.

### Library preparation and sequencing

We used primers that were previously designed to amplify the V3-V4 hyper-variable regions of the 16S rRNA gene (45). A limited cycle PCR generated a single amplicon of ~460 bp, and this was followed by addition of Illumina sequencing adapters and dual-index barcodes. Using paired 300 bp reads, and MiSeq v3 reagents, the ends of each read were overlapped to generate high-quality, full-length reads of the V3 and V4 region in a single run.

### Data Analysis

Sequenced reads were processed for quality assessment, filtering, barcode trimming, and chimera detection were performed on de-multiplexed sequences using the USEARCH method in the Quantitative Insights Into Microbial Ecology (QIIME) package (v.1.9.1) (46). OTUs were defined by clustering with 97% sequence similarity cutoffs (at 3% divergence). The representative sequence for an OTU was chosen as the most abundant sequence showing up in that OTU’s by collapsing identical sequences, and choosing the one that was read the most abundant sequences. Then representative sequences were aligned against Greengenes database core set (v.gg_13_5) using PyNAST alignment method(www.greengenes.secondgenom.com). The minimum sequence length of 150nt and the minimum percent of 75% match were used for the alignment (47, 48). The RDP Classifier program (v.2.2) was used to assign the taxonomy to the representative set of sequences using a pre-built database of assigned sequence of reference set (49). Alpha Diversity was performed using PhyloSeq R package(www.bioconductor.org), Chao1 metric (estimates the species richness.), the observed species metric (the count of unique OTUs found in the sample) and Phylogenetic Distance (PD_whole_tree) were calculated (50). Similarly, Beta Diversity were calculated using QIIME and visualized using Principal Coordinate Analysis (PCoA) to visualize distances between samples on an x-y-z plot. The ranked abundance profile was created using BiodiversityR R Bioconductor package (2.7-2) to highlight the most abundant phylum in all samples. To determine the differentially abundant taxonomic groups over different groups in soman against non-soman exposed at different doses and seizing versus non-seizing groups were examined by fitting linear models using moderated standard errors and the empirical Bayes model following TMM normalization on OTU count. The normalized abundance profile was created across different doses (0.8 and 1.0 LD_50_) of soman exposure compared with control samples, similarly seizing versus non-seizing rats compared against non soman exposed control samples. Sequences can be accesed from the NCBI Sequence Read Archive (SRA) at study accession SRP116704 bioproject PRJNA401162.

To predict the metabolites contributed as per microbial composition we used PICRUSt (v.1.0.0), open source tool that uses precomputed gene content inference for 16S rRNA. PICRSUSt uses the OTU abundance count generated using ‘closed-reference’ OTU picking against Greengenes database, for normalization of OTU table, each OTU was first divided by known as well as predicted 16s copy number abundance (51). Final metagenome functional predictions were performed by multiplying normalized OTU abundance by each predicted functional profile. Statistical hypothesis testing analysis of metagenomics profiles was performed using R to compare KEGG Orthologs (v.80.0) between pre- and post-Soman exposed samples and Principal coordinates analysis was also performed. The predicted metagenome functional counts were normalized using TMM normalization method to fit linear models using contrast function to compute fold changes by moderating the standard errors using empirical Bayes model. Logodds and moderated t-statistic of differential predicted significant metabolite derived from KEGG Orthologs was computed between pre- and post-soman exposed samples at different doses and seizing vs non seizing groups with *p* value cutoff ≤ 0.05. Significant metabolites were further annotated using in-house metabolite annotation function with other databases such as HMDB (v.2.5), KEGG (compounds, pathways, orthologs and reactions) (v.80.0), SMPDB (v.2.0) and FOODB (v.1.0).

### Metabolomic profiling and data analysis

Urine samples were processed using the method of Tyburski (52). Briefly, the samples were thawed on ice and vortexed. For metabolite extraction, 20 μL of urine was mixed with 80 μL of 50% acetonitrile (in water) containing internal standards (10 μL of debrisoquine (1mg/mL) and 50 μL of 4, nitro-benzoic acid (1mg/mL)). The supernatant was transferred to a fresh tube and used for UPLC-ESI-Q-TOF-MS analysis (Xevo G2, Waters Corporation). Each sample (5 μL) was injected onto a reverse-phase 50 × 2.1 mm BEH 1.7 μm C18 column using an Acquity UPLC system (Waters Corporation, USA). The gradient mobile phase was comprised of water containing 0.1% formic acid solution (A) and acetonitrile containing 0.1% formic acid solution (B). Each sample was resolved for 10 min at a flow rate of 0.5 ml/min. This approach has been extensively used for metabolomic profling of biofluids; UPLC gradients conditions and the mass spectrometry parameters and has been described in details (53-55). The UPLC gradient consisted of 100% A for 0.5 min then a ramp of curve 6 to 60% B from 0.5 min to 4.5 min, then a ramp of curve 6 to 100% B from 4.5 to 8.0 min, a hold at 100% B until 9.0 min, then a ramp of curve 6 to 100% A from 9.0 min to 9.2 min, followed by a hold at 100% A until 10 mins. The column eluent was introduced directly into the mass spectrometer by electrospray. Mass spectrometry was performed on a Quadrupole-time-of-flight mass spectrometer operating in either negative or positive electrospray ionization mode with a capillary voltage of 3.2 KV and a sampling cone voltage of 35 V. The desolvation gas flow was 800 L/h and the temperature was set to 350°C. The cone gas flow was 50 L/h, and the source temperature was 150°C. The data was acquired in the V mode with scan time of 0.3 seconds, and inter-scan delay at 0.08 seconds. Accurate mass was maintained by infusing sulfadimethoxine (311.0814 *m/z*) in 50% aqueous acetonitrile (250 pg/μL) at a rate of 30 μL/min via the lockspray interface every 10 seconds. Data were acquired in centroid mode from 50 to 850 *m/z* mass range for TOF-MS scanning, in duplicates (technical replicates) for each sample in positive and negative ionization mode and checked for chromatographic reproducibility. For all profiling experiments, the sample queue was staggered by interspersing samples of the two groups to eliminate bias. Pooled sample injections throughout the run (one pool was created by mixing 2 μL aliquot from all 110 samples) were used as quality controls (QCs) to assess inconsistencies that are particularly evident in large batch acquisitions in terms of retention time drifts and variation in ion intensity over time. QCs were projected in the orthogonal partial least squares-discriminant analysis (OPLS-DA) model along with the study samples to ensure that the technical performance did not impact the biological information(56). The raw data were pre-processed using the XCMS (57) software for peak detection and alignment. The resultant three dimensional data matrix consisting of mass/charge ratios with retention times and feature intensities was subjected to multivariate data analysis using Metaboanalyst v 3.0. Quantitative descriptors of model quality for the OPLS-DA models included R2 (explained variation of the binary outcome: Treatment vs. control and Q2 (cross-validation based predicted variation of the binary outcome). We used score plots to visualize the discriminating properties of the OPLS-DA models. The features selected via OPLS- were used for accurate mass based database search; subsequently the identity of a sub-set of metabolites was confirmed using tandem mass spectrometry.

### Ethics approval and consent to Particpate

This research complied with the Animal Welfare Act and implementing Animal Welfare Regulations, the Public Health Service Policy on Humane Care and Use of Laboratory Animals, and adhered to the principles noted in The Guide for the Care and Use of Laboratory Animals (NRC, 2011).

The views, opinions, and findings contained in this report are those of the authors and should not be construed as official Department of the Army position, policy, or decision, unless so designated by other official documentation. Citations of commercial organizations or trade names in this report do not constitute an official Department of the Army endorsement or approval of the products or services of these organizations.

### Consent for publication

Not Applicable. Human subjects did not participate in this study.

### Availability of data and materials

Data generated and analysed during this study are included in this published aricle and supplementary information files with the exception of raw urine mass spectrometry and 75 day microbiome assay, which will be available from the corresponding author on reasonable request. Sequences can be accesed from the NCBI Sequence Read Archive (SRA) at study accession SRP116704 bioproject PRJNA401162.

## Competing Interest

The authors declare no competing interests. BARDA was not involved in the study design or in the collection, analysis and interpretation of data or the decision to write this manuscript and submit it for publication. The views expressed in this manuscript are those of the authors and do not reflect the official policy of the Department of Army, Department of Defense or the US Government.

## Funding

Support was provided by an interagency agreement between the Biomedical Advanced Research and Development Authority (BARDA), the Geneva Foundation, and the US Army Medical Research Institute of Chemical Defense (USAMRICD) as well as a memorandum of agreement between USAMRICD and US Army Center of Environmental Health (USACEHR). The Metabolomics Shared Resource in Georgetown University (Washington DC, USA) partially supported by NIH/NCI/CCSG grant P30-CA051008.

## Authors’ Contributions

DG analyzed data and wrote the manuscript. AG coordinated microbiome and metabolomics analyses and obtained the samples, AH processed microbiome specimens and completed validation, RK analyzed the data, AKC completed metabolomics analysis. FR analyzed the EEG data, CS conducted animal experiments. LL, MJ and RH conceived and designed the study and edited manuscript. All authors read and approved the final manuscript.

## Acknowledgments

We thank Ms. Kirandeep Gill (Georgetown University) for technical assistance with metabolomics data as well as Dr. Matthew Rice and Dr. Julia Scheerer for editorial input.

